# Single-cell analysis reveals the pan-cancer invasiveness-associated transition of adipose-derived stromal cells into COL11A1-expressing cancer-associated fibroblasts

**DOI:** 10.1101/2020.06.23.166066

**Authors:** Kaiyi Zhu, Lingyi Cai, Chenqian Cui, Dimitris Anastassiou

**Author notes:** Corresponding author: (DA). These authors contributed equally to this work (Kaiyi Zhu, Lingyi Cai).

## Abstract

During the last ten years, many research results have been referring to a particular type of cancer-associated fibroblasts associated with poor prognosis, invasiveness, metastasis and resistance to therapy in multiple cancer types, characterized by a gene expression signature with prominent presence of genes *COL11A1*, *THBS2* and *INHBA*. Identifying the underlying biological mechanisms responsible for their creation may facilitate the discovery of targets for potential pan-cancer therapeutics. Using a novel computational approach for single-cell gene expression data analysis identifying the dominant cell populations in a sequence of samples from patients at various stages, we conclude that these fibroblasts are produced by a pan-cancer cellular transition originating from a particular type of adipose-derived stromal cells naturally present in the stromal vascular fraction of normal adipose tissue, having a characteristic gene expression signature. Focusing on a rich pancreatic cancer dataset, we provide a detailed description of the continuous modification of the gene expression profiles of cells as they transition from *APOD*-expressing adipose-derived stromal cells to *COL11A1*-expressing cancer-associated fibroblasts, identifying the key genes that participate in this transition. These results also provide an explanation to the well-known fact that the adipose microenvironment contributes to cancer progression.

**Author summary:** Computational analysis of rich gene expression data at the single-cell level from cancer biopsies can lead to biological discoveries about the nature of the disease. Using a computational methodology that identifies the gene expression profile of the dominant cell population for a particular cell type in the microenvironment of tumors, we observed that there is a remarkably continuous modification of this profile among patients, corresponding to a cellular transition. Specifically, we found that the starting point of this transition has a unique characteristic signature corresponding to cells that are naturally residing in normal adipose tissue. We also found that the endpoint of the transition has another characteristic signature corresponding to a particular type of cancer-associated fibroblasts with prominent expression of gene *COL11A1*, which has been found strongly associated with invasiveness, metastasis and resistance to therapy in multiple cancer types. Our results provide an explanation to the well-known fact that the adipose tissue contributes to cancer progression, shedding light on the biological mechanism by which tumor cells interact with the adipose microenvironment. We provide a detailed description of the changing profile during the transition, identifying associated genes as potential targets for pan-cancer therapeutics inhibiting the underlying mechanism.

## 1. Introduction

This research is a follow-up of our previous work in 2010 [1], in which we had identified a stage-associated signature of a particular type of cancer-associated fibroblasts (CAFs) by comparing the gene expression of lower stage with that of higher stage cancer samples. The signature, with prominent presence of genes *COL11A1*, *INHBA* and *THBS2*, was identical across cancer types and would only be present after a particular staging threshold, different in each cancer type, was reached. For example, the signature only appeared in ovarian cancer of at least stage III; in colon cancer of at least stage II; and in breast cancer of at least invasive stage I (but never in carcinoma in situ).

We had observed the striking consistency of the signature across cancer types, which was obvious at that time from bulk microarray data. For example, Table 1 shows the top 15 genes ranked in terms of fold change for three different cancer types (breast [2], ovarian [3], pancreatic [4]) using data provided in papers published independently in 2006, 2007, 2008, respectively. The breast cancer data compare invasive ductal carcinoma with ductal carcinoma in situ (supplementary data 3, “up in IDC” of the paper [2]); the ovarian cancer data compare metastatic tissue in the omentum with primary tumor (table 2 of the paper [3]); and the pancreatic data compare whole tumor tissue with normal pancreatic tissue (table 1 of the paper [4]).

**Table 1.** Top 15 ranked genes in terms of fold change (FC) for three different cancer types revealing the signature of the *COL11A1*-expressing cancer-associated fibroblasts. Shown are the rankings, reported by the authors, for breast, ovarian and pancreatic cancers. We eliminated multiple entries of the same gene (keeping the one that appears first) and dashes. Genes shared in all three cancer types are highlighted in green, while genes appearing twice are highlighted in yellow.

|  | Breast |  | Ovarian |  | Pancreatic |  |
| --- | --- | --- | --- | --- | --- | --- |
| Rank | Gene | FC | Gene | FC | Gene | Log2FC |
| 1 | COL11A1 | 6.5 | COL11A1 | 8.23 | INHBA | 5.15 |
| 2 | COL10A1 | 4.07 | COL1A1 | 5.67 | COL10A1 | 5 |
| 3 | MFAP5 | 3.73 | TIMP3 | 5.52 | POSTN | 4.92 |
| 4 | LRRC15 | 3.61 | FN1 | 5.4 | SULF1 | 4.63 |
| 5 | INHBA | 3.44 | INHBA | 4.94 | COL8A1 | 4.6 |
| 6 | FBN1 | 3.43 | EFEMP1 | 4.86 | COL11A1 | 4.4 |
| 7 | SULF1 | 3.35 | DSPG3 | 4.36 | CTHRC1 | 4.38 |
| 8 | GREM1 | 3.35 | COL5A2 | 4.07 | COL1A1 | 4.12 |
| 9 | COL5A2 | 3.22 | LOX | 4.03 | THBS2 | 3.97 |
| 10 | LOX | 3.22 | MFAP5 | 4.01 | HNT | 3.9 |
| 11 | COL5A1 | 3.08 | POSTN | 3.97 | CSPG2 | 3.87 |
| 12 | THBS2 | 2.99 | COL5A1 | 3.95 | WISP1 | 3.8 |
| 13 | LAMB1 | 2.97 | THBS2 | 3.91 | FN1 | 3.69 |
| 14 | FAP | 2.96 | FBN1 | 3.9 | COMP | 3.53 |
| 15 | SPOCK | 2.91 | FAP | 3.84 | COL5A2 | 3.38 |

**Table 2.** Ranked *COL11A1*-associated genes in five PDAC samples.

| Rank | T23 | MI | T11 | MI | T6 | MI | T15 | MI | T18 | MI |
| --- | --- | --- | --- | --- | --- | --- | --- | --- | --- | --- |
| 1 | **COL11A1** | 1 | **COL11A1** | 1 | **COL11A1** | 1 | **COL11A1** | 1 | **COL11A1** | 1 |
| 2 | COL10A1 | 0.3603 | CTHRC1 | 0.2434 | MFAP5 | 0.2353 | MFAP5 | 0.3198 | MFAP5 | 0.3408 |
| 3 | COL12A1 | 0.3383 | MFAP5 | 0.2357 | FNDC1 | 0.1997 | GJB2 | 0.2583 | SUGCT | 0.3379 |
| 4 | COL1A1 | 0.3187 | COL12A1 | 0.2345 | NTM | 0.1912 | COL10A1 | 0.2580 | COL10A1 | 0.2899 |
| 5 | **THBS2** | 0.3167 | COL10A1 | 0.2238 | COL8A1 | 0.1877 | **INHBA** | 0.2561 | C5orf46 | 0.2753 |
| 6 | COL1A2 | 0.3099 | C1QTNF3 | 0.2155 | TWIST1 | 0.1714 | C1QTNF3 | 0.2514 | PPAPDC1A | 0.2668 |
| 7 | COL5A2 | 0.3003 | **THBS2** | 0.2123 | COL10A1 | 0.1619 | MATN3 | 0.2505 | NTM | 0.2649 |
| 8 | CTHRC1 | 0.2854 | COL1A2 | 0.2045 | **THBS2** | 0.1559 | FNDC1 | 0.2503 | COL8A1 | 0.2534 |
| 9 | FN1 | 0.2781 | COL8A1 | 0.2018 | ITGA11 | 0.1556 | COL8A2 | 0.2411 | **INHBA** | 0.2430 |
| 10 | COL3A1 | 0.2770 | AEBP1 | 0.2000 | PPAPDC1A | 0.1305 | COL1A1 | 0.2399 | FNDC1 | 0.2264 |
| 11 | **INHBA** | 0.2746 | LUM | 0.1989 | DIO2 | 0.1298 | COL12A1 | 0.2351 | COL12A1 | 0.2194 |
| 12 | AEBP1 | 0.2688 | COL1A1 | 0.1985 | IGFL2 | 0.1178 | COL8A1 | 0.2325 | IGFL2 | 0.2153 |
| 13 | COL5A1 | 0.2626 | FNDC1 | 0.1963 | SUGCT | 0.1170 | **THBS2** | 0.2292 | **THBS2** | 0.2094 |
| 14 | VCAN | 0.2457 | SFRP2 | 0.1955 | ADAM12 | 0.1165 | NTM | 0.2257 | CTHRC1 | 0.2026 |
| 15 | MFAP5 | 0.2449 | GJB2 | 0.1879 | C1QTNF3 | 0.1165 | COL1A2 | 0.2220 | SULF1 | 0.2015 |
| 16 | MMP11 | 0.2360 | MATN3 | 0.1817 | ITGBL1 | 0.1109 | GREM1 | 0.2156 | COMP | 0.1926 |
| 17 | COL8A1 | 0.2357 | COL3A1 | 0.1740 | GREM1 | 0.1018 | FN1 | 0.2146 | STMN2 | 0.1926 |
| 18 | COL6A3 | 0.2339 | **INHBA** | 0.1696 | P4HA3 | 0.1008 | IGFL2 | 0.2141 | WNT2 | 0.1925 |
| 19 | POSTN | 0.2316 | DCN | 0.1692 | **INHBA** | 0.1002 | CXCL14 | 0.2112 | MMP11 | 0.1919 |
| 20 | MFAP2 | 0.2275 | CTGF | 0.1691 | COL5A1 | 0.0983 | ITGBL1 | 0.2048 | SPOCK1 | 0.1878 |

The four genes *COL11A1*, *INHBA*, *THBS2*, *COL5A2* appear among the top 15 in all three sets (*P* = 6×10^−23^ by hypergeometric test [5]). The actual *P* value is much lower than that, because, in addition to the above overlap, ten additional genes (*COL10A1*, *COL1A1*, *COL5A1*, *FAP*, *FBN1*, *FN1*, *LOX*, *MFAP5*, *POSTN*, *SULF1*) appear among the top 15 in at least two of the three sets (and are highly ranked in all three sets anyway). This similarity demonstrates that the signature is well-defined and associated with a universal biological mechanism in cancer. We had also found that gene *COL11A1* serves as a proxy of the full signature, in the sense that it is the only gene from which all other genes of the signature are consistently top-ranked in terms of the correlation of their expression with that of *COL11A1*. Accordingly, we had identified a *COL11A1*-correlated pan-cancer gene signature, listed in table 4 of [1], which we deposited in GSEA. We had referred to those CAFs as MAFs (“metastasis-associated fibroblasts”), because their presence at the invasive edge of the tumor suggests that metastasis is imminent. To avoid any inaccurate interpretation of the term as implying that such fibroblasts are markers of metastasis that has occurred already, here we refer to them as “*COL11A1*-expressing CAFs.”

Since then, many research results were published connecting one of the genes *COL11A1*, *INHBA*, *THBS2* with poor prognosis, invasiveness, metastasis, or resistance to therapy, in various cancer types [6–15]. Furthermore, several designated tumor subtypes were identified in individual cancer types as a result of the presence of those pan-cancer CAFs. For example, the top 15 genes distinguishing the ovarian “mesenchymal subtype” according to [16] are *POSTN*, *COL11A1*, *THBS2*, *COL5A2*, *ASPN*, *FAP*, *MMP13*, *VCAN*, *LUM*, *COL10A1*, *CTSK*, *COMP*, *CXCL14*, *FABP4*, *INHBA*. Similarly, the 24 characterizing genes of the “activated stroma subtype” of pancreatic cancer in fig. 2 of [17] are *SPARC*, *COL1A2*, *COL3A1*, *POSTN*, *COL5A2*, *COL1A1*, *THBS2*, *FN1*, *COL10A1*, *COL5A1*, *SFRP2*, *CDH11*, *CTHRC1*, *PNDC1*, *SULF1*, *FAP*, *LUM*, *COL11A1*, *ITGA11*, *MMP11*, *INHBA*, *VCAN, GREM1*, *COMP*. In both cases, the subtype-defining signature is clearly the *COL11A1*/*INHBA*/*THBS2*-expressing CAF signature.

In this work we show that the origin of those *COL11A1*-expressing CAFs is a type of adipose-derived stromal/stem cells (ASCs) naturally present in the stromal vascular fraction (SVF) of normal adipose tissue, expressing a unique characteristic signature containing fibroblastic markers such as *LUM* and *DCN* as well as adipose-related genes, such as *APOD*, *CFD* and *MGP*. This finding explains the stage association of the *COL11A1*-expressing signature as resulting from the interaction of tumor cells with the adipose microenvironment: Indeed, adipose tissue is encountered when ovarian cancer cells reach the omentum (stage III); after colon cancer has grown outside the colon (stage II); and in breast cancer from the beginning of the spread (stage I, but not in situ stage 0).

Our main computational methodology is attractor analysis (Materials and Methods), and is used in a novel manner for the analysis of rich single-cell RNA sequencing (scRNA-seq) data. The unsupervised attractor algorithm [18] iteratively finds co-expression signatures converging to “attractor metagenes” pointing to the core (“heart”) of co-expression. Each attractor metagene is defined by a ranked set of genes along with scores determining their corresponding strengths within the signature, so the top-ranked genes are the most representative of the signature. The attractor algorithm has previously been used successfully for identifying features useful for breast cancer prognosis [19,20]. When applied on single cell data from a sample, it identifies the gene expression profiles of the dominant cell populations in the sample. Accordingly, we demonstrate that there is a continuous “ASC to *COL11A1*-expressing CAF transition.”

We used publicly available scRNA-seq data sets from five different cancer types. We first focused on one of those datasets [21] from pancreatic ductal adenocarcinoma, because it is exceptionally rich, containing gene expression profiles from 57,530 cells in 24 tumor samples (T1-T24) and 11 normal control samples (N1-N11). Using attractor analysis, we established that (a) the dominant fibroblastic population in each of the 11 normal samples is identical to that of the ASC type described above, and (b) that the gene expression profiles of the dominant fibroblastic population of the 24 tumor samples undergo a gradual change as the transition proceeds, starting from the state of the normal ASCs and passing through a continuum of intermediate states. We also validated these results by applying trajectory inference analysis. Finally, we validated our results in the other cancer types (head and neck, ovarian, lung, breast), suggesting the pan-cancer nature of the ASC to *COL11A1*-expressing CAF transition.

## 2. Results

### 2.1 Pancreatic Ductal Adenocarcinoma (PDAC) dataset

To find the expression profile of the dominant fibroblastic population in each sample, we applied the attractor algorithm on the set of identified mesenchymal cells (Materials and Methods). We used the general fibroblastic marker gene *LUM* as seed, and found that all samples yielded strong co-expression signatures involving many genes with big overlap among them. S1 Table shows the top 100 genes for each of the 34 samples (11 normal and 23 tumor samples, excluding sample T20 as it did not contain identified fibroblasts). Genes *LUM*, *DCN*, *FBLN1*, *MMP2*, *SFRP2* and *COL1A2* appear in at least 33 out of the 34 samples, revealing a strong similarity shared by all those fibroblastic expression profiles. This strong overlap is consistent with the continuous transition process, as described below.

#### 2.1.1 Establishing the presence of *COL11A1*-expressing CAFs

Because it serves as proxy of the full signature [1], a reliable test for determining if a sample contains the *COL11A1*-expressing CAFs is to rank all genes in terms of their association with *COL11A1* and see if *INHBA* and *THBS2* are top ranked. Indeed, this happens in several tumor samples, as shown in Table 2 for some of them (T23, T11, T6, T15, T18). For each sample, the shown genes are co-expressed in the same cells, because of the high correlations in a single-cell dataset.

#### 2.1.2 Dominant fibroblastic populations in the tumor samples

Table 3 shows the ranked lists of genes for the attractors in each of the rearranged tumor samples. There is a remarkable continuity in the shown expression profiles. The samples at the right side of the table include *COL11A1* at increasingly higher ranks. On the other hand, we identified gene *APOD* (Apolipoprotein D) as particularly significant, as it is highly ranked on the left side and its rank gradually decreases as we go to the right. The intermediate tumor samples shown in the middle have cells expressing genes that are top-ranked in both the lists on the left as well as on the right. In other words, these cells are in a genuine intermediate state, rather than being a mixture of distinct subtypes. Furthermore, all genes shown in Table 3 have high strengths and therefore are co-expressed inside the same individual cells in each sample.

**Table 3.**
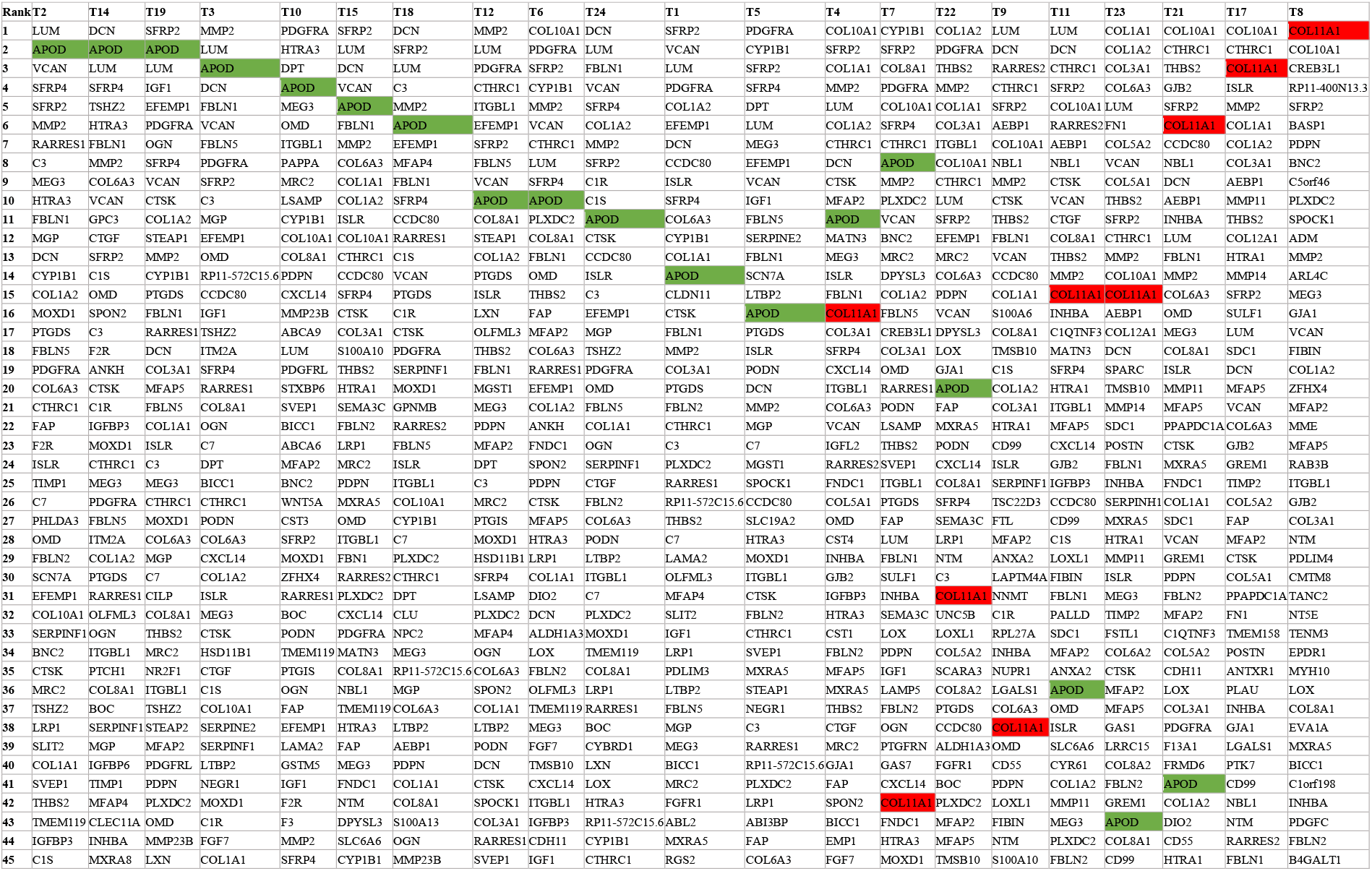
Rearranged PDAC tumor samples showing the continuously changing pattern of the signature profile. Genes *APOD* and *COL11A1* are highlighted in green and red, respectively.

#### 2.1.3 Dominant fibroblastic population in the normal samples

Table 4 shows the attractor metagenes of the 11 normal samples by listing the 20 top-ranked genes for each sample, in which commonly shared genes are color-coded. There is a striking similarity among them, indicating that they represent a stable and normally occurring cell population. Specifically, there are 18 genes shared among the top 50 genes in the attractors of at least ten of the eleven normal samples, listed by average rank (*P* < 10^−7^ by permutation test). In addition to fibroblastic markers, there are several strongly expressed adipose-related or stemness-related genes in the list, such as *APOD*, *CFD*, *CXCL12* and *DPT*, revealing that they are ASCs. Consistently, Gene Set Enrichment Analysis of these 18 genes identified most significant enrichment (FDR *q* value = 5.38×10^−29^) in the “BOQUEST_STEM_CELL_UP” dataset of genes upregulated in stromal stem cells from adipose tissue versus the non-stem counterparts [22].

**Table 4.**
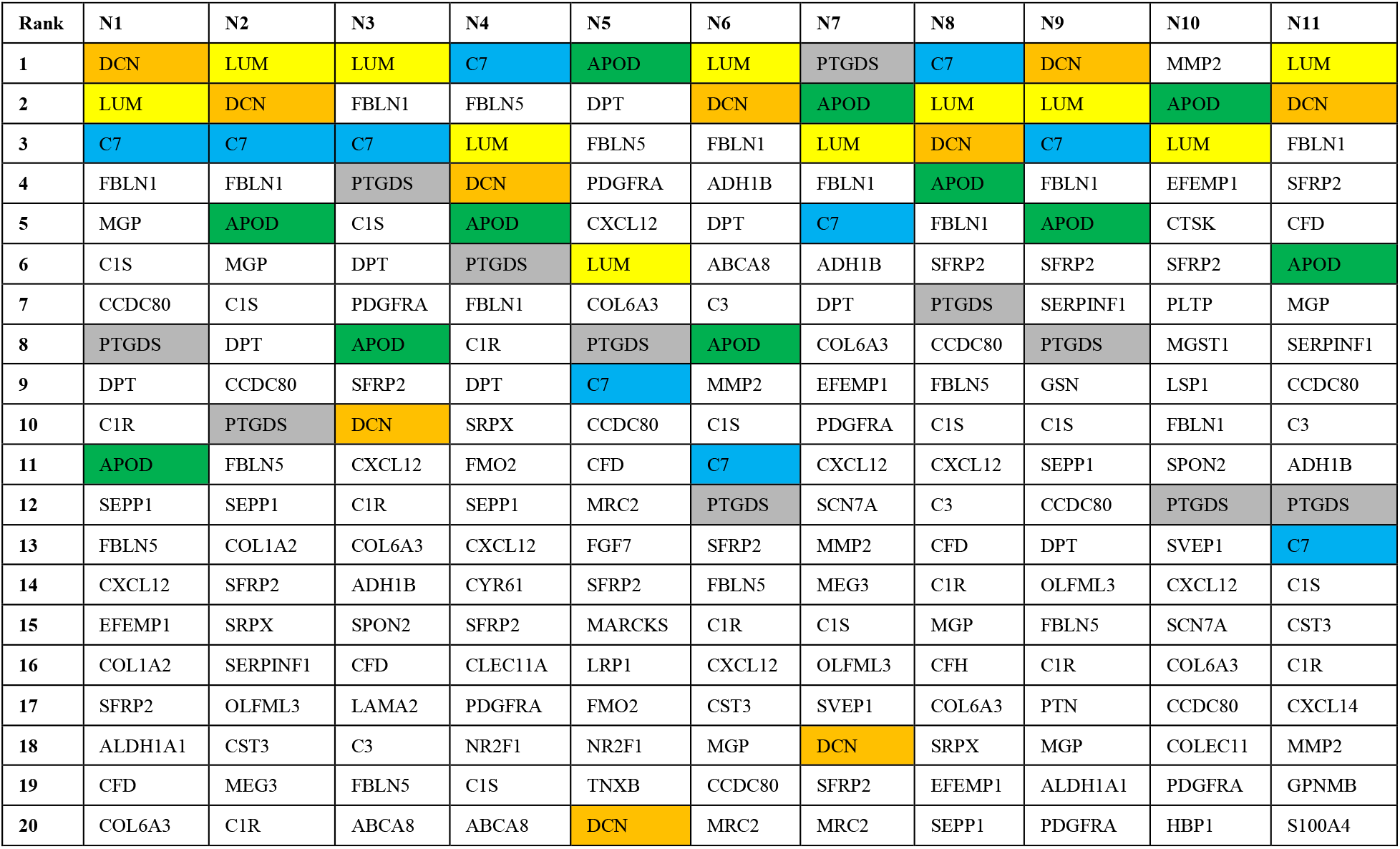
Top 20 genes of the identified attractors for each pancreatic normal sample (N1-N11).

To investigate the nature of this ASC population, we referred to recent results from single-cell analysis of general human adipose tissue [23]. We used the dataset with the single-cell expression profiles of all 26,350 cells taken from the SVF of normal adipose tissue from 25 samples. The first column in Table 5 contains the top 30 ranked genes in the corresponding attractor signature. The second column contains the top 30 ranked genes of the consensus attractor (Materials and Methods) of the 11 normal pancreatic samples, representing the main state of the normal fibroblastic population. There are 14 overlapping genes in the two lists (*P* = 10^−37^), and most of the non-highlighted genes in each column are still ranked highly in the other column. This extreme similarity of the two gene expression profiles indicates that they correspond to the same naturally occurring cell population.

**Table 5.**
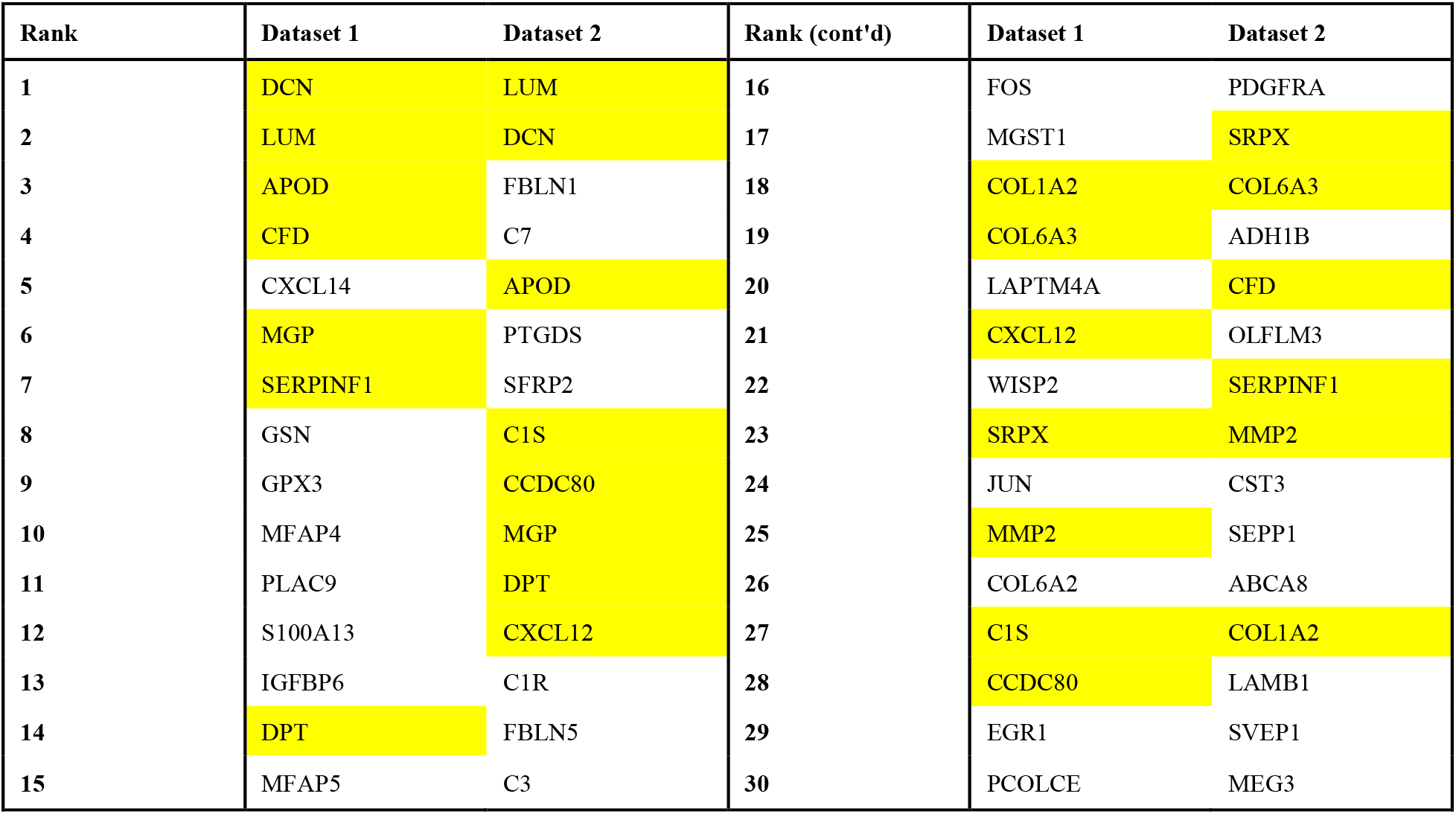
Comparison of the attractors (top 30 genes) identified in the SVF of normal adipose tissue (Dataset 1) and in the normal pancreatic samples (Dataset 2). Common genes are highlighted in yellow.

| Rank | Dataset 1 | Dataset 2 | Rank (cont'd) | Dataset 1 | Dataset 2 |
| --- | --- | --- | --- | --- | --- |
| 1 | DCN | LUM | 16 | FOS | PDGFRA |
| 2 | LUM | DCN | 17 | MGST1 | SRPX |
| 3 | APOD | FBLN1 | 18 | COL1A2 | COL6A3 |
| 4 | CFD | C7 | 19 | COL6A3 | ADH1B |
| 5 | CXCL14 | APOD | 20 | LAPTM4A | CFD |
| 6 | MGP | PTGDS | 21 | CXCL12 | OLFLM3 |
| 7 | SERPINF1 | SFRP2 | 22 | WISP2 | SERPINF1 |
| 8 | GSN | C1S | 23 | SRPX | MMP2 |
| 9 | GPX3 | CCDC80 | 24 | JUN | CST3 |
| 10 | MFAP4 | MGP | 25 | MMP2 | SEPP1 |
| 11 | PLAC9 | DPT | 26 | COL6A2 | ABCA8 |
| 12 | S100A13 | CXCL12 | 27 | C1S | COL1A2 |
| 13 | IGFBP6 | C1R | 28 | CCDC80 | LAMB1 |
| 14 | DPT | FBLN5 | 29 | EGR1 | SVEP1 |
| 15 | MFAP5 | C3 | 30 | PCOLCE | MEG3 |

#### 2.1.4 Further demonstration of the continuity of the transition

As an additional confirmation of the continuity of the transition (as opposed to the presence of a mixture of distinct fibroblastic subtypes), Fig 1 shows scatter plots, color-coded for the expression of fibroblastic marker *LUM*, for the fibroblast-rich samples T11 and T23, demonstrating the presence in each sample of cells representing the full range of the continuously varying transition from *APOD* to *COL11A1*-expressing cells.

**Fig 1.**
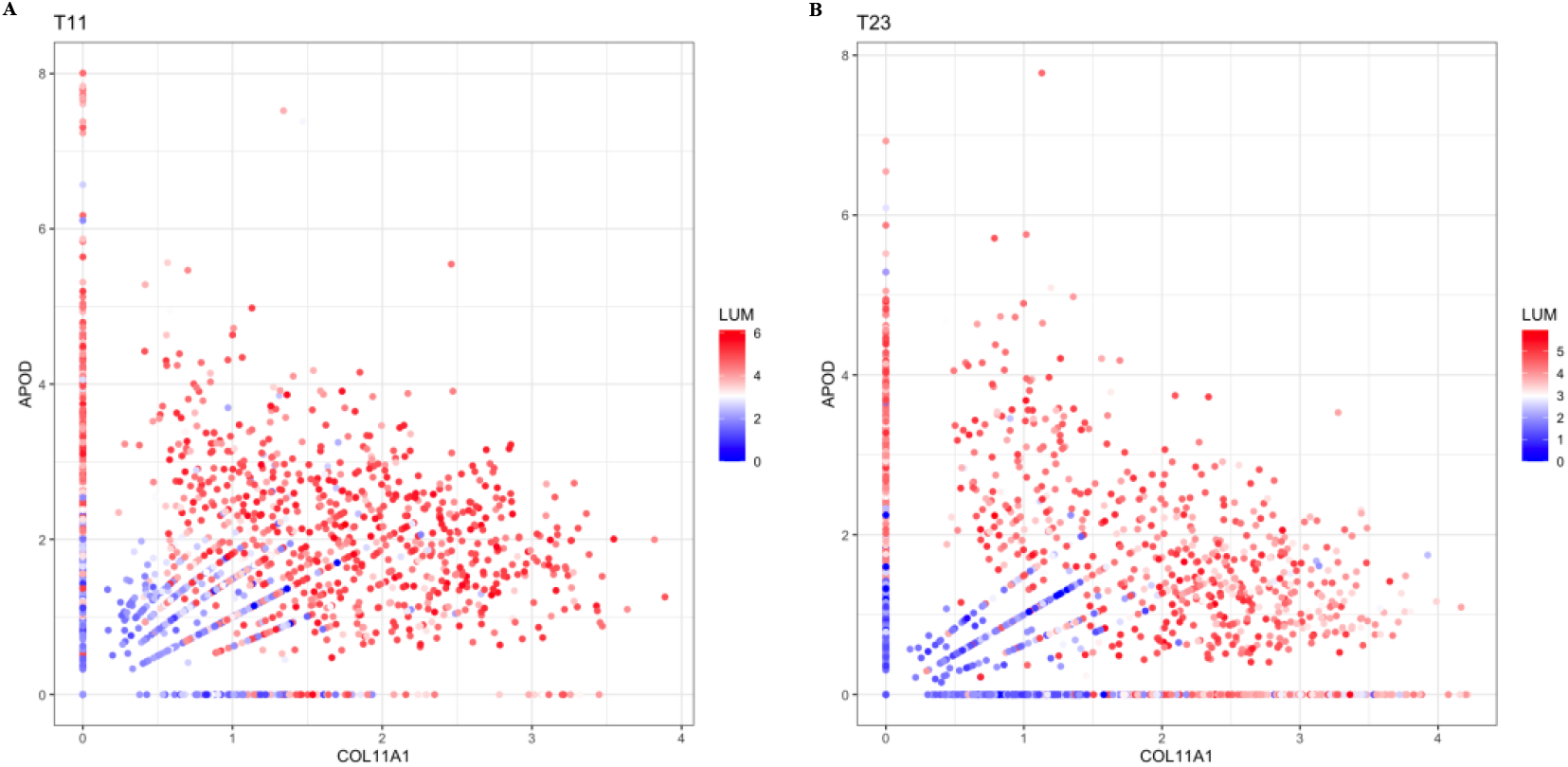
Scatter plots for fibroblast-rich samples for patients T11 and T23.

To further investigate the continuous transition, we partitioned the 34 pancreatic samples into three groups, considering whether the corresponding attractor includes *COL11A1* among the top 100 genes (S1 Table). Group 1 includes the eleven normal samples (N1 to N11). Group 2 includes twelve tumor samples in which *COL11A1* is not included (T1, T2, T3, T5, T10, T12, T13, T14, T16, T18, T19, T24). Group 3 includes eleven cancer samples in which it is included (T4, T6, T7, T8, T9, T11, T15, T17, T21, T22, T23). We then applied the consensus version of the attractor finding algorithm (Materials and Methods) and identified the signatures representing the main state of the fibroblasts for each of the above three sample groups (Table 6), so we can observe the genes whose ranking changes significantly during the transition.

**Table 6.** Top 50 genes of the consensus attractors for three different PDAC sample groups. Group1: normal samples; Group 2: tumor samples in which the dominant fibroblastic population does not include *COL11A1*; Group 3: tumor samples in which the dominant fibroblastic population includes *COL11A1*.

| Rank | Group1 | Group2 | Group3 | Rank (cont'd) | Group1 | Group2 | Group3 |
| --- | --- | --- | --- | --- | --- | --- | --- |
| 1 | LUM | LUM | COL1A1 | 26 | ABCA8 | PLXDC2 | NTM |
| 2 | DCN | DCN | COL10A1 | 27 | COL1A2 | COL1A1 | NBL1 |
| 3 | FBLN1 | APOD | COL1A2 | 28 | LAMB1 | OGN | ITGBL1 |
| 4 | C7 | SFRP4 | CTHRC1 | 29 | SVEP1 | C1S | FN1 |
| 5 | APOD | SFRP2 | SFRP2 | 30 | MEG3 | PTGDS | MFAP2 |
| 6 | PTGDS | VCAN | COL3A1 | 31 | PCOLCE | ITGBL1 | RARRES2 |
| 7 | SFRP2 | FBLN1 | MMP2 | 32 | NEGR1 | TSHZ2 | GJB2 |
| 8 | C1S | MMP2 | LUM | 33 | FGF7 | PODN | MXRA5 |
| 9 | CCDC80 | PDGFRA | VCAN | 34 | LRP1 | SERPINF1 | LOX |
| 10 | MGP | EFEMP1 | THBS2 | 35 | CYR61 | OMD | MEG3 |
| 11 | DPT | RARRES1 | COL6A3 | 36 | ALDH1A1 | COL10A1 | MRC2 |
| 12 | CXCL12 | FBLN5 | DCN | 37 | EFEMP1 | HSD11B1 | HTRA1 |
| 13 | C1R | ISLR | COL11A1 | 38 | FMO2 | IGF1 | PDPN |
| 14 | FBLN5 | COL6A3 | APOD | 39 | CFH | RP11-572C15.6 | FBLN2 |
| 15 | C3 | C3 | SFRP4 | 40 | PLTP | LTBP2 | C1QTNF3 |
| 16 | PDGFRA | CCDC80 | COL5A2 | 41 | SLIT2 | FBLN2 | TMSB10 |
| 17 | SRPX | CTSK | FAP | 42 | MRC2 | COL3A1 | POSTN |
| 18 | COL6A3 | CTHRC1 | FBLN1 | 43 | SCN7A | STEAP1 | CDH11 |
| 19 | ADH1B | CYP1B1 | COL8A1 | 44 | LAMA2 | LXN | MMP14 |
| 20 | CFD | MEG3 | COL5A1 | 45 | TIMP1 | BOC | CCDC80 |
| 21 | OLFML3 | COL1A2 | AEBP1 | 46 | COL6A2 | MRC2 | SULF1 |
| 22 | SERPINF1 | MOXD1 | MFAP5 | 47 | CXCL14 | LRP1 | TMEM119 |
| 23 | MMP2 | C7 | INHBA | 48 | LTBP4 | THBS2 | CXCL14 |
| 24 | CST3 | MGP | CTSK | 49 | EMP1 | MFAP4 | FNDC1 |
| 25 | SEPP1 | COL8A1 | ISLR | 50 | GSN | FAP | IGFBP3 |

Although there are many shared genes, the groups have distinct gene rankings. Group 1 (normal samples) contains many adipose-related genes, consistent with Table 4. Group 3 contains, in addition to *COL11A1*, many among the other CAF genes, such as *THBS2*, *INHBA*, *MMP11*, *AEBP1*, *MFAP5*, *COL10A1* and *VCAN*.

Group 2 displays an intermediate state, including markers of both ASCs as well as CAFs. To find the most unique and representative gene(s) for Group 2, we focused on the top 20 genes of its attractor and calculated the ranking increment when compared with the rankings of the same genes in the attractors of the other two groups. We considered a ranking increase of greater than 50 as significant. As a result, we found that *RARRES1* is the only gene satisfying the criterion in both comparisons.

Gene *RARRES1* (aka *TIG1*) appears in the beginning of the ASC to *COL11A1*-expressing CAF transition process and in all samples of Group 2. This is consistent with the known fact that it plays an important, though not yet clear, role in regulating proliferation and differentiation of adipose tissue derived mesenchymal stem cells [24]. On the other hand, the expression of *RARRES1* is reduced in Group 3, disappearing completely in the final stages of differentiation when *COL11A1* is expressed strongly (consistent with suggestions that it is a tumor suppressor [25,26]).

We also performed differential expression (DE) analysis on the classified fibroblasts between the three sample groups defined above (Materials and Methods; see complete DE gene lists in S2 Table). Importantly, the results of such analysis represent the full population of fibroblasts and not necessarily the expression changes in the particular cells undergoing the ASC to *COL11A1*-expressing CAF transition.

For the ‘Group1 to Group 2’ stage, genes *CFD* and *DPT* are the most downregulated genes, consistent with the downregulation of adipose-related genes. On the other hand, the top upregulated gene is phospholipase A2 group IIA (*PLA2G2A*), whose encoded protein is a member of a family of enzymes catalyzing the hydrolysis of phospholipids into free fatty acid. At the same time, other extracellular matrix genes are also strongly upregulated, as expected.

For the ‘Group 2 to Group 3’ stage, remarkably *PLA2G2A* again shows as the top DE gene, however this time it is down-regulated. Similarly, the tumor suppressor *RARRES1*, which was significantly up-regulated in the early stage becomes down-regulated in this stage. On the other hand, the most significantly upregulated genes, following the top gene *COL11A1*, were matrix metalloproteinase (*MMP11*), collagens of type X and XII (*COL10A1* and *COL12A1*), thrombospondin 2 (*THBS2*), growth factor binding protein 3 (IGFBP3), while *CFD*, *APOD* and *C7* continued decreasing their expression levels. Thus, this stage represents the final formation of *COL11A1*-expressing CAFs.

The top differentially expressed gene *PLA2G2A* is not among the top genes of any attractors we identified and is expressed by less than half of cells, even in Group 2 in which it appears. Therefore, it is expressed by cells in Group 2 that are not among the ones undergoing the ASC to *COL11A1*-expressing CAF transition, although it probably still plays an important related role and many previous studies referred to its effects on prognosis of multiple cancer types [27–29]. This gene is a phospholipase catalyzing the hydrolysis of phospholipids into fatty acids. On the other hand, it has been recognized that fatty acid oxidation is associated with the final *COL11A1*-expressing stage of the transition [30]. These results suggest that lipid metabolic reprogramming plays an important role in the metastasis-associated biological mechanism [31], providing energy for the metastasizing tumor cells.

#### 2.1.5 Trajectory Inference

We independently applied trajectory inference analysis on the PDAC fibroblasts by using the Slingshot [32] method. We first performed unsupervised clustering on the identified fibroblasts (Materials and Methods), resulting in four subgroups X1, X2, X3, X4 (S1A Fig) with the top differentially expressed genes shown in S1B Fig. One of these clusters (X4) was discarded from further TI analysis, because it mainly expressed the IL1 CAF marker *HAS1* (Hyaluronan Synthase 1), which is not expressed by either ASCs or *COL11A1*-expressing CAFs (and does not appear at all in S1 Table), and contained only 3% of fibroblasts resulting almost exclusively from patient T11 (S1C Fig).

As seen from the list of top differentially expressed genes, X1 contains CAF genes top ranked (including *MMP11*, *COL11A1*, *THBS2*, *INHBA*), X2 has *RARRES1* at the top, and X3 has ASC genes top ranked, including *DPT*, *C7*, *CXCL12* and *CFD*. Consistently, S2A Fig and S2B Fig show the single trajectory path resulting from TI analysis, where X3 is the starting point and X1 is the end point of the trajectory, while X2 (highly expressing *RARRES1*), is an intermediate point, thus validating the continuous ASC to *COL11A1*-expressing CAF transition. S3 Table shows the top 100 genes with zero *P* value, ranked by their variances, resulting from pseudotime-based differential gene expression analysis (Materials and Methods). We can clearly identify as top-ranked several ASC genes, as well as CAF genes, while some general fibroblastic markers, such as *DCN*, are missing, consistent with the continuity of the ASC to *COL11A1*-expressing CAF transition. We then used a generalized additive model (GAM) fit to pseudotime-ordered expression data to visualize the trend of gene expressions (Fig 2A).

**Fig. 2.**
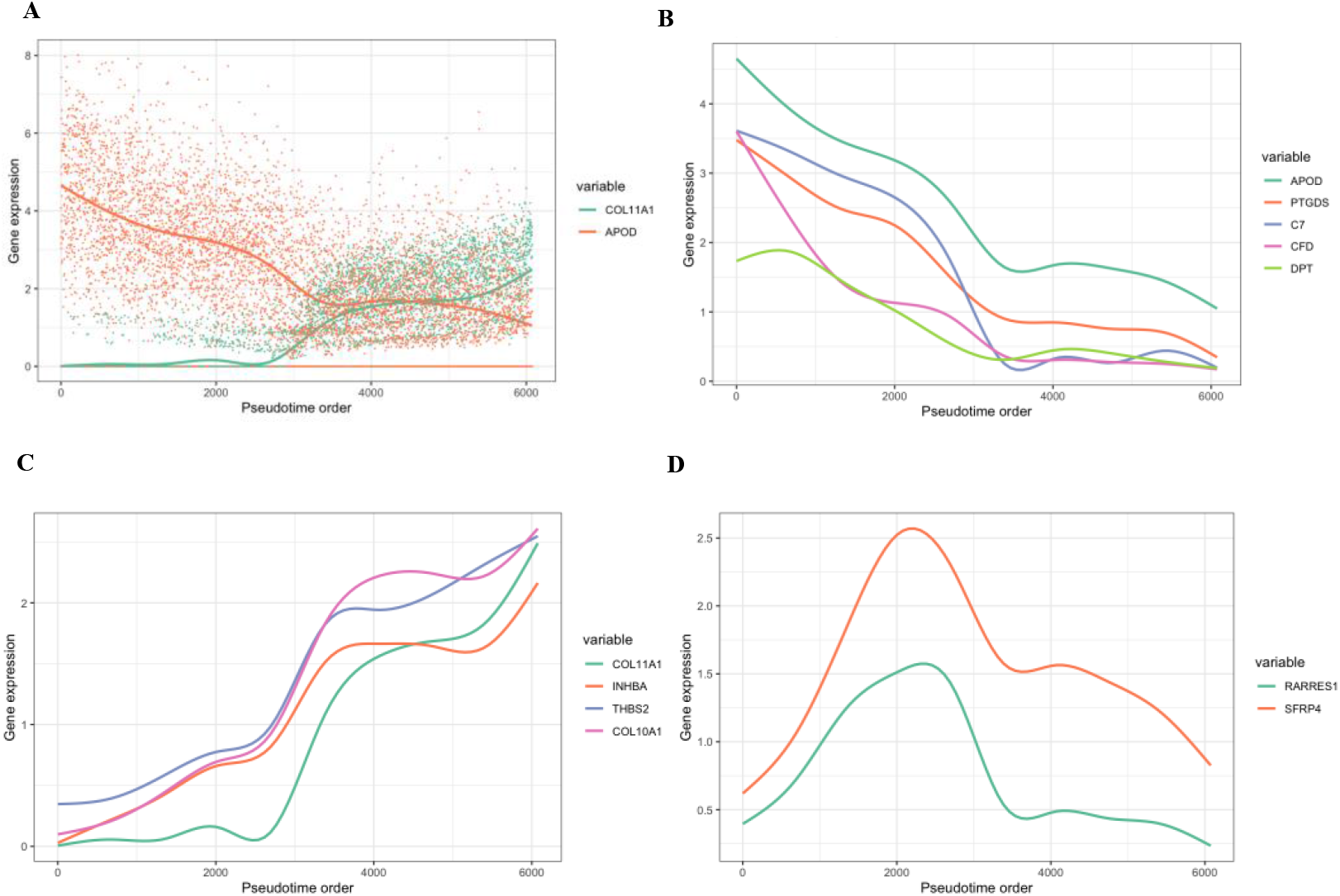
Trajectory analysis of PDAC. **A.** GAM fit to pseudotime ordered expression data to visualize the trend of gene expressions. **B.** Expression of adipose-related genes along the transition lineage. The x axis shows the cell orders and the y axis shows the normalized read count. **C.** Expression of *COL11A1*-associated genes along the transition lineage. **D.** Expression of *RARRES1* and *SFRP4* genes along the transition lineage.

There was a prominent difference between adipose-related genes and *COL11A1*-associated genes. The expression of the adipose-related genes steadily fell across the process (Fig 2B), while the expression of *COL11A1*-associated genes gradually increased (Fig 2C). There is a significant negative correlation between these two groups of genes, e.g., *COL11A1* (the last among those genes to increase its expression) was exclusively overexpressed in the mature CAFs, which did not express *C7*. Of particular interest, genes *RARRES1* and *SFRP4* (Fig 2D) increased consistently at the beginning and then decreased after reaching a peak, suggesting that they play important roles in the differentiation path.

### 2.2 Validation in other cancer types

Next, we validated the ASC to *COL11A1*-expressing CAF transition in other solid cancer types. Although we could not find currently available datasets as rich as the PDAC dataset, we selected those containing a large (at least 100) number of fibroblasts and separately analyzed each of them, obtaining consistent results. Specifically, we used four scRNA-seq datasets from head and neck cancer (HNSCC) [33], ovarian cancer [34], lung cancer [35] and breast cancer [36].

The *COL11A1*-expressing CAF signature has been confirmed to be a pan-cancer signature [37–39]. Therefore, the most important validation task would be to confirm the existence of the *APOD*/*CFD*/*CXCL12*/*MGP*/*PTGDS*-expressing ASCs as the starting point of the transition, and to also confirm that some samples are at an intermediate stage, expressing genes such as *RARRES1*, *SFRP4* and *THBS2*, in addition to the core ASC genes, demonstrating that they are at an intermediate stage of the transition.

#### 2.2.1 Head and neck squamous cell carcinoma

For the HNSCC dataset, the authors of the paper presenting the data [33] reported that the cancer-associated fibroblasts in the dataset can be partitioned into two subsets, which they name CAF1 and CAF2. In table s5 of that paper, the top three differentially expressed genes of the CAF2 group are *CFD*, *APOD* and *CXCL12*, while the full gene list for CAF2 presented in the same table s5 also includes genes *MGP*, *C3*, *C7*, *DPT*, *PTGDS*. This strongly suggests that the partitioning used in the paper was influenced by the presence of an ASC cell subpopulation, identical, or at least very similar to, those discovered in the PDAC. Similarly, the list of differentially expressed genes for CAF1 in table s5 includes genes *INHBA*, *THBS2*, *CTHRC1*, *POSTN*, *MMP11*, *COL5A2*, *COL12A1*, suggesting that the identified CAF1 subpopulation was influenced by the presence of differentiated CAFs, which would eventually express *COL11A1*. Finally, gene *RARRES1* also appears among the list of CAF2 genes, suggesting that it was captured among cells which had started the process of ASC to *COL11A1*-expressing CAF transition.

In our independent analysis, we performed clustering identifying 1,026 fibroblasts from all available cells (S3A Fig; Materials and Methods). There were two fibroblastic clusters (X7 and X9) expressing CAF associated genes (*COL11A1*, *COL12A1*, *MMP11*, *INHBA*, *THBS2*, *COL10A1*, *COL8A1*, *FN1*) and ASC associated genes (*APOD*, *C7*, *PTGDS*), respectively (S4 Table), which confirmed the presence of these two populations in HNSCC.

Among the individual patients, we found that the most prominent case is sample HNSCC28, which contains a rich set of cells undergoing differentiation. Applying the attractor finding algorithm on the fibroblasts of that sample (S5 Table) resulted in genes *LUM*, *APOD*, *COL6A3*, *PDGFRA*, *DCN*, and *CFD* being among the top-ranked, revealing that it represents an ASC population. Furthermore, the presence of genes *THBS2*, *MFAP5* and *VCAN* in the same attractor reveals that these cells have already started undergoing the transition.

#### 2.2.2 Ovarian cancer

For the ovarian dataset, the clustering results showed two clusters (X6 and X9) expressing *COL11A1*-associated genes and ASC-associated genes, respectively (S3B Fig, S6 Table; Materials and Methods). Among the individual patients, we found that the ones validating our hypotheses most are HG2F and LG2, both of whose datasets, consistently, contain cells from the fatty omental tissue. S5 Table includes the corresponding two attractors identified in the cells of each patient. Among the top ranked genes for HG2F are *DCN*, *LUM*, *C1S*, *C7*, and *C3*, but also *RARRES1*, suggesting that they represent fibroblasts undergoing the transition, while the LG2-based attractor contains highly ranked all three genes *COL11A1*, *INHBA*, *THBS2*.

#### 2.2.3 Lung cancer

The dataset contains a large number (> 50,000) of cells, but we only classified ~2% (= 1,346) among them as mesenchymal cells, including fibroblasts and pericytes (Materials and Methods). Among those cells, there were two fibroblastic clusters (X1 and X2) expressing related genes (*COL11A1*, *COL12A1*, *MMP11*, *INHBA*, *THBS2*, *COL10A1*, *COL8A1*, *FN1*) and ASC related genes (*APOD*, *C7*, *PTGDS*), respectively (S3C Fig, S7 Table). The presence of the transition is evident by the attractors identified in the mesenchymal cells for patients 4 and 3 (S5 Table). The former prominently contains genes *CFD*, *PTGDS* and *C7*, while the latter contains *THBS2*, *COL10A1* and *INHBA*.

#### 2.2.4 Breast cancer

The size of the breast cancer dataset is small (~1,500 cells in total), and 169 cells among them were classified as mesenchymal (Materials and Methods). By further clustering these cells, we identified ASCs (X1) and *COL11A1*-expressing CAFs (X3) (S3D Fig, S8 Table). ASC related genes (*APOD*, *MFAP4*, *CFD*) were identified in X1, while CAF-related genes (*COL10A1*, *COL11A1*, *MMP11*, *INHBA*, *FN1*, *THBS2*, *AEBP1*, *COL12A1*) are among the top 15 of X3. Patients PT089 and PT039 contain the highest proportions (>50%) of the ASC and *COL11A1*-expressing CAF subpopulations, respectively, and we found consistent results in their attractors (S5 Table), as the former contains *C1S*, *C1R*, *CXCL12*, *PTGDS*, *C3*, while the latter contains *THBS2*, *COL11A1*, *COL10A1*, at top-ranked positions.

### 2.3 Potential therapeutic targets

This work provides opportunities for identifying therapeutic targets inhibiting the cellular transition. For example, targeting of gene *MFAP5* was recently found to enhance chemosensitivity in ovarian and pancreatic cancers [40]. Specifically, the author states that “*MFAP5* blockade suppresses fibrosis through downregulating of fibrosis-related genes such as *COL11A1*.” Consistently, we found *MFAP5* as one of the most highly associated genes with *COL11A1* (Table 2).

As mentioned earlier, genes *SFRP4* and *RARRES1* are transiently expressed in Group 2 of Table 6, suggesting that they can be investigated for inhibiting the cellular transition. Consistently, gene *RARRES1* (aka *TIG1*) was found to play an important, though not yet clear, role in regulating proliferation and differentiation of adipose tissue derived mesenchymal stem cells [24], while *SFRP4* is a Wnt pathway regulator whose expression has been found associated with various cancer types [41,42].

Of particular interest as potential drivers are noncoding RNAs due to their typical regulatory role. Because the expression of these genes is not accurately captured by scRNA-seq technology, we did a thorough analysis of the full set of The Cancer Genome Atlas (TCGA) pan-cancer data. For the RNA sequencing and miRNA sequencing dataset of each cancer type, we removed the genes in which more than 50% of the samples have zero counts. Then we performed quantile normalization using the limma package [43] (v3.40.6) on log2 transformed counts. In each of the 33 cancer types, we ranked all protein-coding genes in terms of the association (using the metric of mutual information) of their expression with that of gene *COL11A1*. We excluded the 11 cancer types (*LGG*, *SKCM*, *SARC*, *LAML*, *PCPG*, *GBM*, *TGCT*, *THYM*, *ACC*, *UVM*, *UCS*) in which neither *THBS2* nor *INHBA* was among the 50 top-ranked genes, because of the absence of significant amounts of *COL11A1*-expressing CAFs in those samples (1st sheet in S9 Table). In each of the remaining 22 cancer types, we then ranked all long noncoding RNAs (lncRNAs) and microRNAs (miRNAs) in terms of their association with *COL11A1* (2nd and 3rd sheets in S9 Table). Finally, we did pan-cancer sorting of all lncRNAs and miRNAs in terms of the median rank of all lncRNAs and miRNAs (4th sheet in S9 Table).

We found that *LINC01614* represents a particularly promising therapeutic target. It had a perfect score of 1 in the pan-cancer sorting list, being strikingly top-ranked (top1) in 14 (*BRCA*, *UCEC*, *KIRC*, *HNSC*, *LUAD*, *LUAD*, *LUSC*, *OV*, *STAD*, *ESCA*, *PAAD*, *MESO*, *DLBC*, *CHOL*) out of the 22 cancer types (2nd sheet in S9 Table). In fact, the association of *LINC01614* was even higher than that of marker protein-coding gene *INHBA*. We also found that the three top-ranked miRNAs were *miR-199a-1*, *miR-199b*, *miR-199a-2*. The associated *miR-214* is also very highly ranked (3rd sheet in S9 Table).

## 3. Discussion

One contribution of our work is that it provides an explanation to the fact that adipose tissue contributes to the development and progression of cancer as a result of its interaction with malignant cells [44–46].

Another contribution is that it sheds light on one important mechanism within the heterogeneous tumor microenvironment, which also contains fibroblasts that do not participate in the transition. There are several published results, often conflicting with each other, identifying various distinct subtypes of fibroblasts, resulting from computational clustering methods following dimensionality reduction, or using particular marker genes (such as *αSMA*, which, interestingly we found that is not consistently expressed in the *COL11A1*-expressing CAFs, therefore it should not be used as its marker), which represent an approximation of biological reality. Examples include the hC1 and hC0 clusters in [47], the C9 and C10 clusters in [39], the CAF2 and CAF1 clusters in [33], the iCAF and myCAF clusters in [48,49] and the iCAF an mCAF clusters in [50]. A review of such results in pancreatic cancer appears in [51].

As an example of conflicting results, the “iCAFs” identified in [50] are identical to the normal ASCs (fig. 3b of [50]) as evidenced by the list of its differentially expressed genes (*PTGDS*, *LUM*, *CFD*, *FBLN1*, *APOD*, *DCN*, *CXCL14*, *SFRP2*, *MMP2*, all of which appear in Table 5, further validating the ASC signature. Therefore, this identified cluster contains mainly normal cells at the origin of the transition, which should not even be called CAFs. On the other hand, the iCAFs identified in [49] also have significant overlap in their marker composition with the ASCs (such as *CFD*, *DPT* and *CXCL12*), while omitting essential ASC markers such as *APOD*, and including additional genes.

On the other hand, our work is consistent with, and complementary to the results of [47] focusing on the immunotherapy response, in which the presence of the “TGF-beta CAFs” was inferred by an 11-gene signature consisting of *MMP11*, *COL11A1*, *C1QTNF3*, *CTHRC1*, *COL12A1*, *COL10A1*, *COL5A2*, *THBS2*, *AEBP1*, *LRRC15*, *ITGA11*. This population apparently represents the *COL11A1*-expressing CAF endpoint of the transition, and gene *LRRC15* was selected as the representative gene based on the fact that it was found to be the most differentially expressed gene between CAFs and normal tissue fibroblasts in mouse models. Indeed, *LRRC15* is a key member of the *COL11A1*-expressing CAF signature (table 4 of [1]) and we also found that *COL11A1* is the highest associated gene to *LRRC15* in the Group 3 PDAC patients,

In our work we used a detailed gene association-based scrutiny of all our results, including numerous color-coded scatter plots, observing and confirming the presence of cells expressing particular combinations of genes, rather than blindly accepting clustering results. We believe that this nontraditional computational methodology, when used on rich single-cell data, represents a paradigm shift in which systems biology alone can be trusted, by itself, for producing reliable results. We hope that our results will give rise to testable hypotheses that could eventually lead to the development of pan-cancer therapeutics inhibiting the ASC to *COL11A1*-expressing CAF transition.

## 4. Materials and methods

### 4.1 Datasets availability

The pancreatic dataset [21] was downloaded from the Genome Sequence Archive with accession number CRA001160. The four validation datasets of other cancer types are also publicly available: HNSCC [33] (GSE103322), ovarian [34] (GSE118828), lung cancer [35] (E-MTAB-6149 and E-MTAB-6653), breast cancer [36] (GSE118389). We excluded samples from lymph nodes. The numbers of patients included in these datasets are 35 (PDAC), 18 (HNSCC), 9 (ovarian), 5 (lung), and 6 (breast).

### 4.2 Data processing and cell identification

We applied the Seurat (v3.1.4) [52] R package to process the gene expression matrix and characterize the cell type identity for each scRNA-seq dataset. The count matrix was normalized and log transformed by using the NormalizeData function. We selected the 2,000 most variable features and then performed principal component analysis (PCA) followed by applying an unsupervised graph-based clustering approach. We used default parameter settings in all the above steps except that the resolution parameter in the FindCluster function is set to 1.0 to increase the granularity of downstream clustering. To identify differentially expressed genes for each cluster, we used the FindMarkers function. To characterize the identity of mesenchymal cells in each dataset, we made use of the expression of known markers: *LUM*, *DCN*, *COL1A1* for fibroblasts, and *RGS5* for pericytes.

For the smaller-size datasets (ovarian, breast), we performed clustering once on all cells for mesenchymal cell identification. For datasets of larger size (PDAC, HNSCC, lung), we applied ‘two-step clustering’ to ensure accuracy: The first step was initial clustering within individual samples. Then we excluded samples with very few (< 20) detected fibroblasts and pooled the mesenchymal cells of the remaining samples together for a second clustering, which resulted in the final set of mesenchymal cells for the dataset. For the PDAC dataset, we included an additional step to remove low-quality cells, by retaining cells for which at least one of the corresponding markers had expression levels ≥ 3.

### 4.3 Mutual information

Mutual information (MI) is a general measure of the association between two random variables [53]. We used a spline based estimator [54] to estimate MI values and normalized so the maximum possible value is 1. The details of the estimation method are described in the paper introducing the attractor algorithm [18]. We used the getMI or getAllMIWz function implemented in the cafr R package with parameter negateMI = TRUE.

### 4.4 Attractor-based analysis

For single dataset, we applied the attractor finding algorithm using the findAttractor function implemented in the cafr (v0.312) R package [18] with gene *LUM* as seed. The exponent (*a*) was set to different values for scRNA-seq datasets profiled from different protocols. For the analysis of UMI based (e.g. 10x) and full-length-based (e.g. Smart-seq2) datasets, we used *a* = 3 and *a* = 5, respectively. Briefly, the attractor finding algorithm iteratively finds mutually associated genes from an expression matrix, converging to the core of the co-expression mechanism. The association measure used is the normalized mutual information (as described above), which captures the general relationships (including nonlinear effects) between variables. To find the consensus attractor for multiple datasets, we applied the consensus version of the attractor finding algorithm as described in [55]. In the consensus version, the association measures between genes are evaluated as the weighted median of the corresponding measures taken from the individual datasets. The weights are proportional to the number of samples included in each individual dataset in log scale.

### 4.5 Trajectory inference (TI) analysis

We selected the Slingshot [32] method for TI analysis, based on its robustness and suggestions made by the benchmarking pipeline dynverse [56]. We used the raw counts as input and followed the Slingshot lineage analysis workflow (v1.4.0). To begin this process, Slingshot chose robustly expressed genes if it has at least 10 cells that have at least 1 read for each. After gene filtering, we proceeded to full quantile normalization. Following diffusion map dimensionality reduction, Gaussian mixture modelling was performed to classify cells. The final step of lineage inference analysis used the slingshot wrapper function. A cluster based minimum spanning tree was subjected to describe the lineage. After analyzing the global lineage structure, we fitted a generalized additive model (GAM) for pseudotime and computed *P* values. Genes were ranked by *P* values and variances. After running Slingshot, we identified genes whose expression values significantly vary over the derived pseudotime by using a GAM, allowing us to detect non-linear patterns in gene expression.

### 4.6 Statistical analysis

#### 4.6.1 P value evaluation by permutation test

The significance of the consistency of N different attractors of size M was evaluated as follows. We randomly selected M genes out of the 24,005 genes to generate a random gene set, and we created N such random gene sets. Each time, we counted the number of genes common in all N gene sets, and we repeated this process ten million times. In these ten million experiments, it never occurred that there is one or more gene common in all N gene sets. Therefore, the corresponding *P* value is less than 10^−7^.

#### 4.6.2 Differential expression analysis

We used a Wilcoxon Rank Sum test by applying the FindMarkers function in Seurat to identify the differentially expressed (DE) genes between fibroblasts of different groups. DE genes with |log fold change| > 0.25 and Bonferroni adjusted *P* value < 0.1 are considered as significant. The positive and negative DE genes are ranked separately in terms of the absolute values of their log fold-change.

## Acknowledgments

The noncoding RNA-related results presented here are in whole based upon data generated by the TCGA Research Network: https://www.cancer.gov/tcga.

## Supplementary Material

**S1 Fig. Overview of the PDAC fibroblasts. A.** 6,267 fibroblasts originated from 11 control pancreases and 23 tumor samples were petitioned into four groups X1-X4. Fractions of the fibroblasts were: 45%, 38%, 14%, and 3%. **B.** Table showing the top 20 DE genes for each cluster. **C.** Bar plots presenting the numbers of cells captured for each cluster.

**S2 Fig. Trajectory analysis of 6,075 fibroblasts in PDAC dataset. A.** Colors coded for pseudotime changing, red presenting the beginning of differentiation and blue presenting the end. **B.** Color-coded trajectory analysis of fibroblasts for annotated three clusters.

**S3 Fig. Unsupervised clustering of four datasets from HNSCC, ovarian cancer, lung cancer and breast cancer. A.** t-SNE embedding of the whole HNSCC dataset. **B.** t-SNE embedding of the whole ovarian cancer dataset. **C.** t-SNE embedding of the mesenchymal cells from lung cancer dataset. **D.** t-SNE embedding of the mesenchymal cells from breast cancer dataset.

**S1 Table. *LUM*-seeded attractors (top 100 genes) identified in each PDAC sample.**

**S2 Table. Differentially expressed genes of between the fibroblasts from different PDAC sample groups.**

**S3 Table. Top 100 genes of temporally expressed genes on the pseudotime variable.**

**S4 Table. Differentially expressed genes among different clusters of HNSCC dataset.**

**S5 Table. *LUM*-seeded attractors (top 100 genes) from validating samples of other cancer types.**

**S6 Table. Differentially expressed genes among different clusters of ovarian cancer dataset.**

**S7 Table. Differentially expressed genes among different clusters of mesenchymal cells from lung cancer dataset.**

**S8 Table. Differentially expressed genes among different clusters of stromal cells from breast cancer dataset.**

**S9 Table. Ranked genes lists in terms of their association (mutual information) with gene *COL11A* by using the full set of pan-cancer TCGA bulk RNA-seq data.**

