## Supplementary material for "Single-cell analysis reveals the pan-cancer invasiveness-associated transition of adipose-derived stromal cells into COL11A1-expressing cancer-associated fibroblasts": S1 Figure

(a)

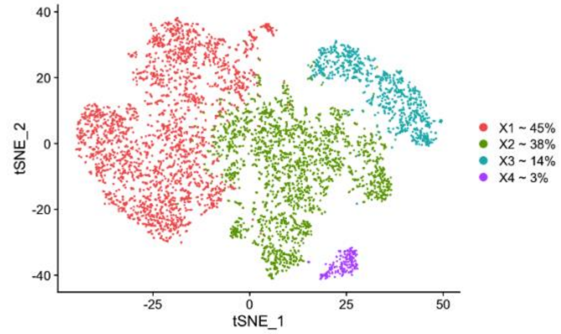


(b)

| | X1 | X2 | X3 | X4 |
|---|---|---|---|---|
| 1 | MMP11 | RARRES1 | PRSS1 | HAS1 |
| 2 | COL11A1 | MGP | DPT | AREG |
| 3 | FN1 | TIMP1 | CLPS | HAS2 |
| 4 | C1QTNF3 | COL14A1 | CTRB1 | NR4A3 |
| 5 | CTHRC1 | C1R | PTN | IL6 |
| 6 | COL10A1 | CST3 | C7 | SEMA6A |
| 7 | GJB2 | C7 | ADH1B | RP1-3J17.3 |
| 8 | COL12A1 | SOD3 | FXYD2 | GPRC5A |
| 9 | SDC1 | MFAP4 | PNLIP | KLF4 |
| 10 | POSTN | CTSC | SEPP1 | TFPI2 |
| 11 | COL1A1 | C1S | AMY2A | NR4A2 |
| 12 | IGFL2 | SERPINF1 | FMO2 | CREM |
| 13 | COL5A2 | IGFBP7 | INS | KDM6B |
| 14 | THBS2 | C3 | CXCL12 | PLAUR |
| 15 | MMP14 | SFRP4 | SYCN | ADAMTS4 |
| 16 | INHBA | SPARCL1 | CFD | SAT1 |
| 17 | COL1A2 | TGM2 | ALDH1A1 | MLLT11 |
| 18 | GREM1 | F2R | CPA1 | MEDAG |
| 19 | PLAU | TSHZ2 | FBLN5 | NFATC2 |
| 20 | CD55 | OGN | PTGDS | TUBB2A |

(c)

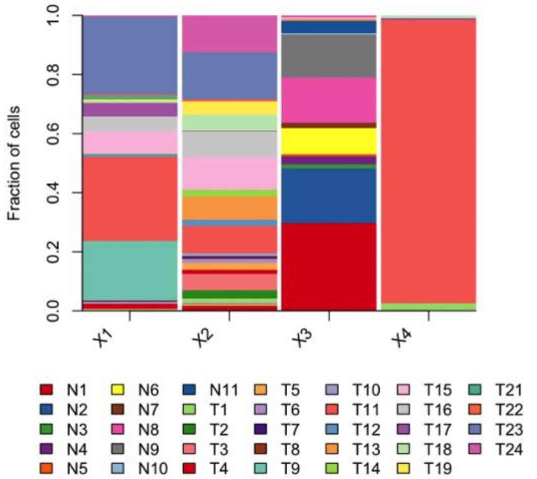
