## Supplementary figures and images for "Single-cell analysis reveals the pan-cancer invasiveness-associated transition of adipose-derived stromal cells into COL11A1-expressing cancer-associated fibroblasts"

### S2 Figure

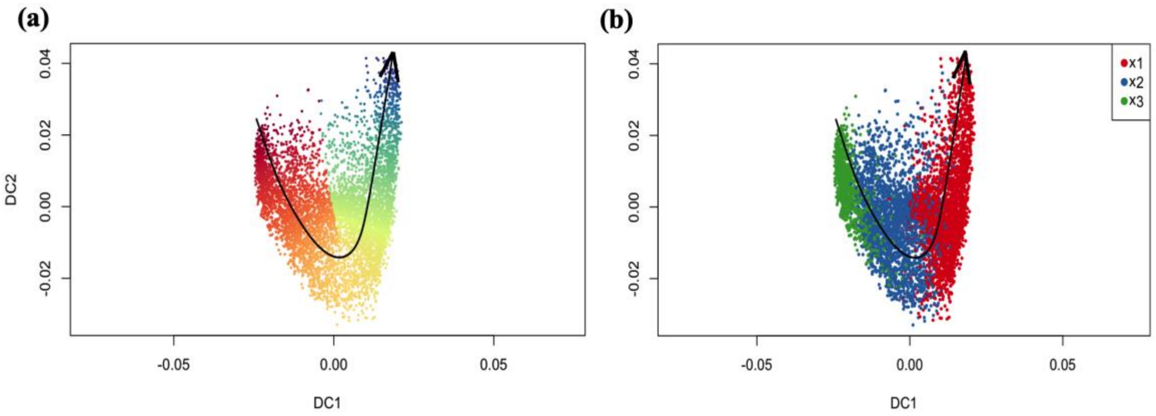

### S3 Figure

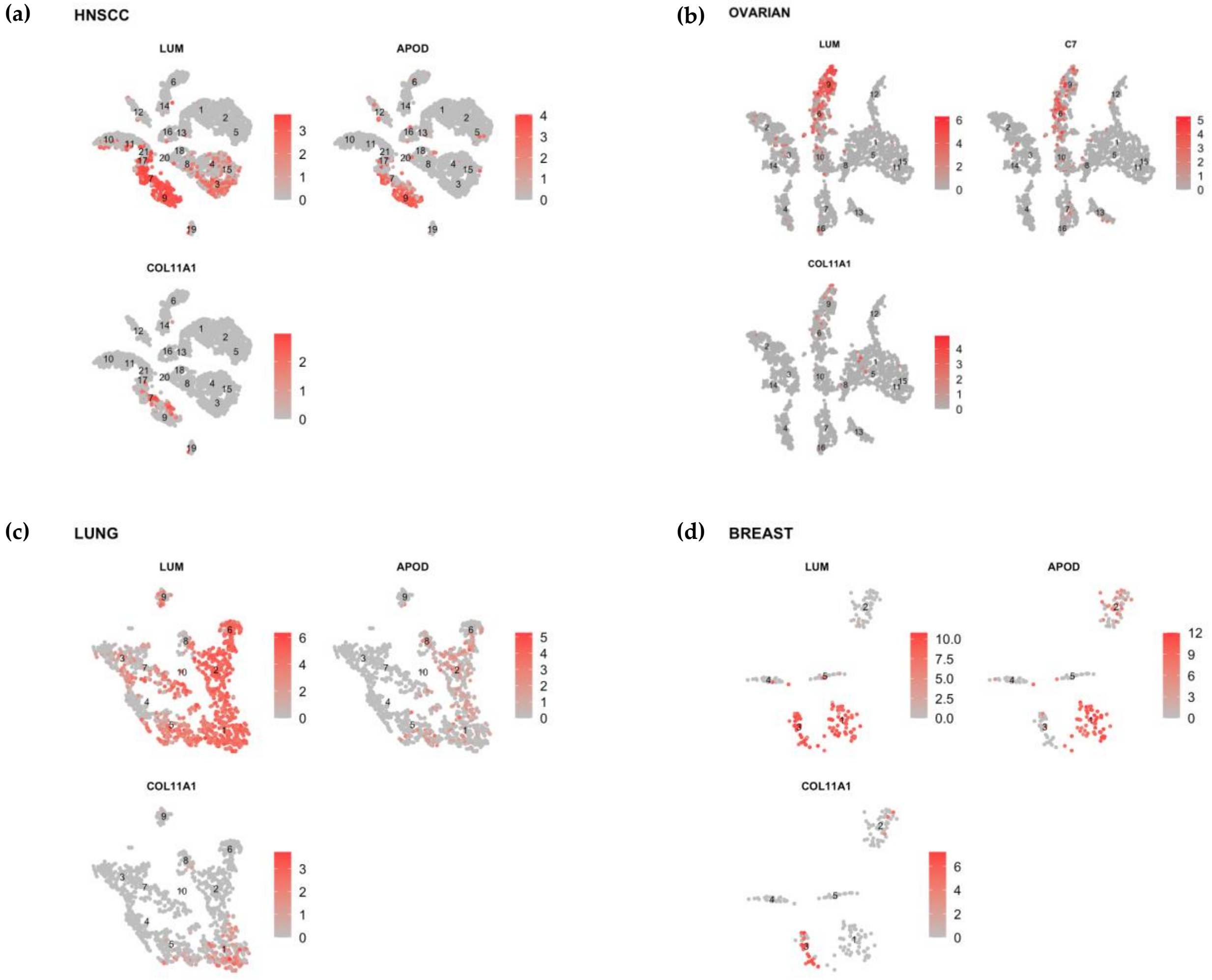
